# A *de novo* reference genome of the red seaweed *Furcellaria lumbricalis*

**DOI:** 10.64898/2026.11.17.767248

**Authors:** Louka Noyen, Antoine Van de Vloet, Frederik Leliaert, Sofie D’hondt, Jordi Morcillo, Tim De Pooter, Mojca Strazisar, Jessica Knoop, Kris Audenaert, Olivier De Clerck

## Abstract

*Furcellaria lumbricalis* is a red seaweed that occurs across a broad salinity gradient, ranging from the saline waters of the NE Atlantic to the brackish Baltic Sea. In the Baltic Sea, natural populations of *F. lumbricalis* are harvested commercially, primarily for *furcellaran* extraction, a sulphated polysaccharide used as a stabilizing, thickening, and gelling agent. Genomic resources for this species remain absent, constraining evolutionary, ecological, and applied research. Here, we present a *de novo* reference genome assembly of the seaweed *Furcellaria lumbricalis* using PacBio Hifi and Oxford Nanopore Technologies long-read sequencing. The final assembly, consisting of 149 contigs spanning 918.2 Mb, contains a high proportion of repetitive elements (90.5%), and has an N50 of 15,5 Mb. We further integrated RNA-seq data, protein homology evidence, and *ab initio* gene prediction to build a genome annotation comprising 6100 genes, of which 82.6% has a functional annotation. This genomic evidence contributes to a steadily growing body of genomic resources of red algae, with often expanded and highly repetitive genomes, and helps facilitate the study of genetic adaptation to their unique environments.

## Background and summary

To date, Rhodophyta (red algae) remain largely overlooked at the genomic level compared to other multicellular eukaryotic lineages, limiting comparative analyses and efforts to understand the basis of their complex multicellularity, physiological adaptation and unique triphasic life cycle^1–3^. Despite the rapidly evolving sequencing technologies, the generation of high-quality genomic resources for these seaweeds remains challenging due to high concentrations of complex sulphated polysaccharides and polyphenols, which interfere with DNA extraction and impair the performance of long-read sequencing technologies^2,4,5^. Although chromosome-level genome assemblies have recently become available for several economically important red seaweeds, including species of the genera *Gracilaria*^6,7^, *Porphyra*^8^, and *Pyropia*^9–11^, genomic representation across the Rhodophyta remains extremely limited, considering that the group is estimated to comprise between 10,000 and 20,000 species^1^^2^. The currently available red algal genomes are generally considered to be relatively compact, with most assemblies ranging from several tens to a few hundred megabases in size^2,6,13^. However, the recent chromosome-level assembly of *Bostrychia moritziana* revealed a genome size exceeding 1 Gb^14^, questioning the longstanding observation that red algal genomes are relatively small and intron-poor and supporting the hypothesis of substantial genome expansion events within the Florideophyceae^2,6,14,15^.

In addition to their largely unexplored genomic characteristics, Rhodophyta are considered as one of the richest sources of biologically active metabolites with applications in food, feed, medical, and biotechnological industries^16–18^. In seaweeds, high concentrations of secondary metabolites are often linked to their occupation of the intertidal zone, where they are exposed to multiple environmental stressors, including tidal desiccation, UV-radiation, salinity fluctuations, and hydrodynamic stress^19,20^. Besides the intertidal, most Rhodophyta specialize in inhabiting the deeper zones, enabled by the presence of specialized accessory photosynthetic pigments, phycobiliproteins, that efficiently absorb blue-green wavelengths penetrating deeper into the water column^21^. This specialized light-harvesting capability produces a distinct ecological niche and reduces direct competition with other macroalgal groups, such as Chlorophyta (green algae) and Phaeophyta (brown algae), which lack these red pigments and therefore predominantly occupy the shallower intertidal and subtidal zones^21,22^.

The red alga *Furcellaria lumbricalis* (hereafter referred to as *Furcellaria*), also known as Clawed Fork Weed, exemplifies strong ecological adaptation by inhabiting an exceptionally broad ecological range, extending from the saline waters of the NE Atlantic to the brackish waters of the Baltic Sea, accompanied by a pronounced morphological variation across its distribution (Figure 1)^23^. In the Baltic Sea, it represents one of the few red seaweeds capable of persisting under low-salinity conditions (down to 4 PSU)^24^, a remarkable degree of osmotic tolerance within the red lineage^25–27^. This physiological characteristic has generated increasing interest in the extraction of bioactive metabolites and osmoprotective compounds with potential applications in agriculture and biotechnology, particularly in the context of abiotic stress mitigation^28–30^.

**Figure 1.**
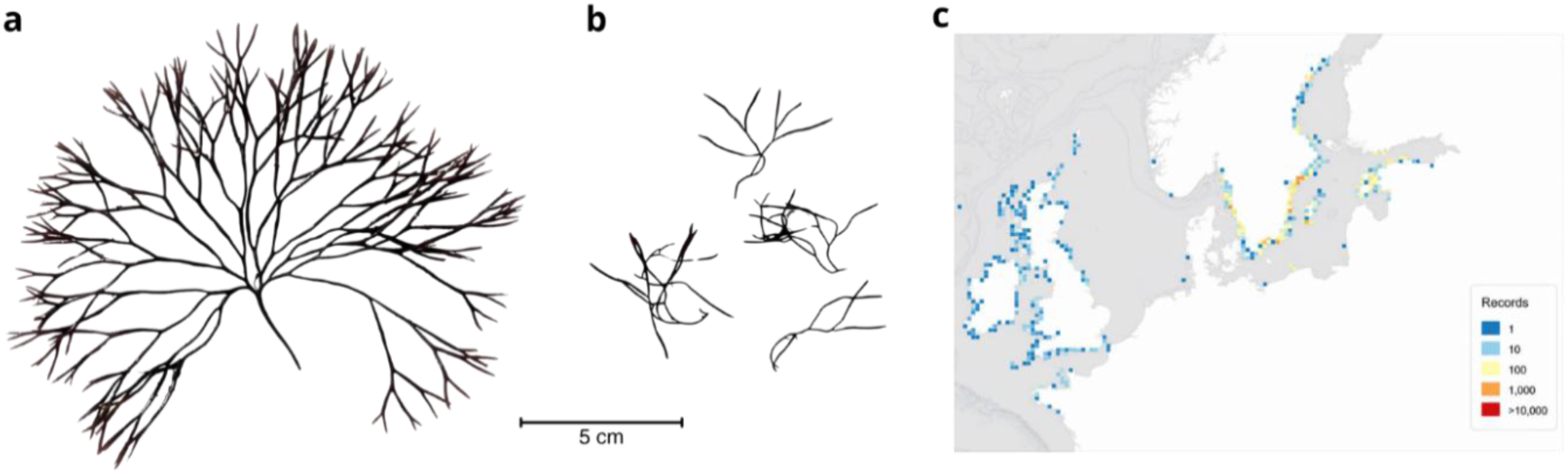
Morphological differences between two ecotypes of *Furcellaria* **a)** thallus of Atlantic (Roscoff, France) *Furcellaria*, **b)** thallus of Baltic (Saaremaa, Estonia) *Furcellaria*, **c)** European distribution map of *Furcellaria* (OBIS^33^).

At present, commercial exploitation of *Furcellaria* relies solely on the harvesting of wild stocks^17^. Consequently, genomic resources are essential to improve our understanding of population structure, to identify signals of local (mal)adaptation, to predict future distributional responses to environmental change, and to facilitate the development of tailored cultivation and management strategies^31,32^.

Here, we present the first reference genome assembly of *Furcellaria* generated through hybrid integration of PacBio HiFi and Oxford Nanopore Technologies (ONT) long-read sequencing data. HiFi sequencing provided high-accuracy long reads, whereas ONT sequencing generated substantially greater long-read coverage. The integration of both datasets improved assembly contiguity and completeness for the highly repetitive *Furcellaria* genome. The final assembly spans 918,200,210 bp and consists of 149 contigs, with the largest contig measuring 44.3 Mb. Additional assembly statistics are summarized in Table 1. BUSCO assessment using the eukaryota_odb10 (74.9%) and rhodophyta_odb12 (95.5%) datasets indicated high completeness within the reported red algal genome assemblies to date (Figure 2). Repeat annotation revealed that 90.5% of the genome consists of repetitive elements. Gene prediction, through combined integration of RNA-seq and homologous protein evidence, identified 6100 protein-coding genes of which 4509 (82.6%) were functionally annotated. Although this first assembly is neither chromosome-level nor telomere-to-telomere, it does provide the necessary resolution for applications in population genomics and adaptation assays.

**Figure 2.**
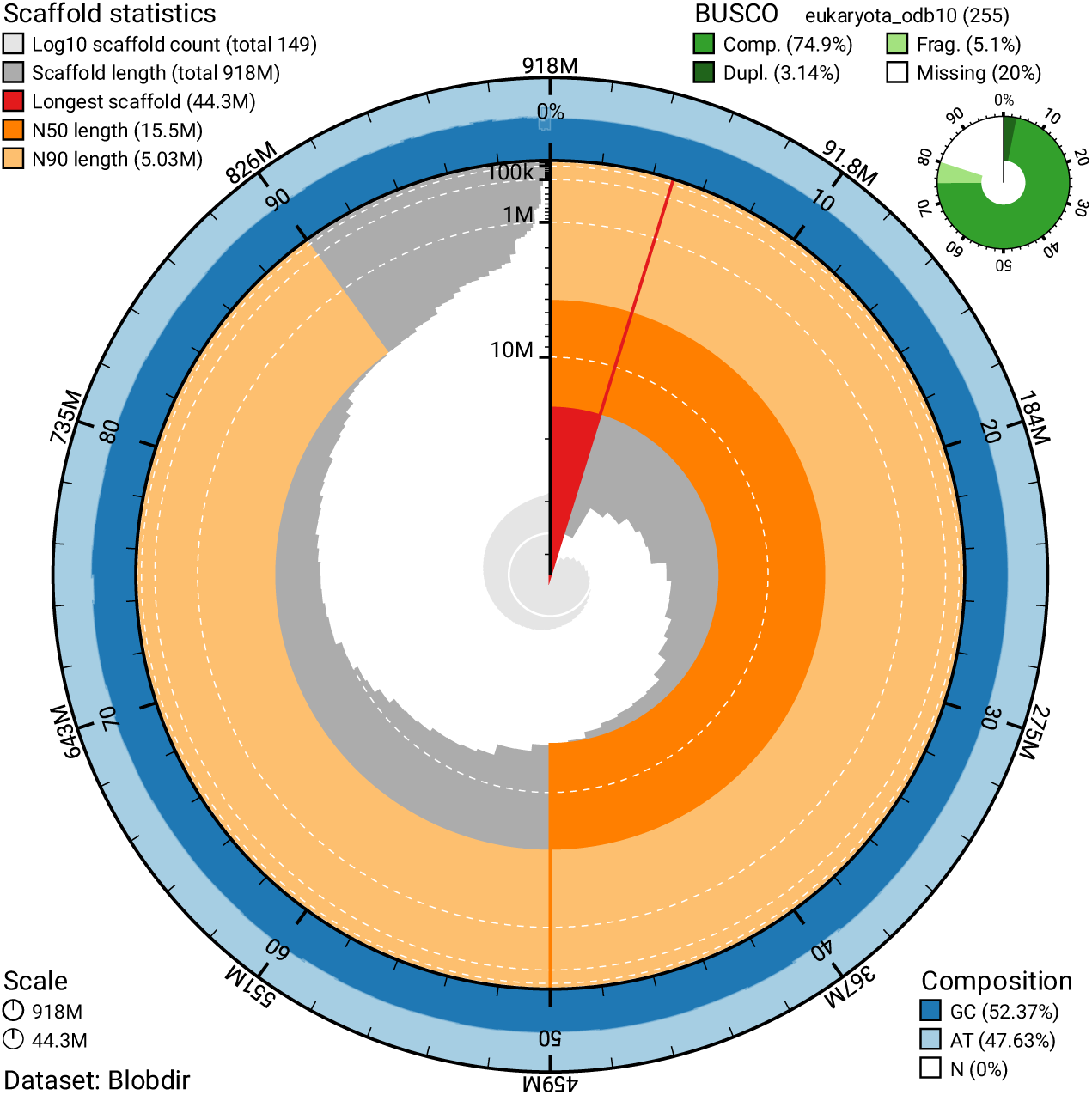
Snail plot of the *Furcellaria* genome assembly, indicating scaffold statistics and BUSCO completeness.

**Table 1.** QUAST summary of the *Furcellaria* genome assembly in terms of number and dimension of its defined contigs.

| **Assembly** | **Number** |
|---|---|
| # contigs (>= 0 bp) | 149 |
| # contigs (>= 1000 bp) | 149 |
| # contigs (>= 5000 bp) | 149 |
| # contigs (>= 10000 bp) | 148 |
| # contigs (>= 25000 bp) | 142 |
| # contigs (>= 50000 bp) | 112 |
| Total length (>= 0 bp) | 918,200,210 |
| Total length (>= 1000 bp) | 918,200,210 |
| Total length (>= 5000 bp) | 918,200,210 |
| Total length (>= 10000 bp) | 918,194,252 |
| Total length (>= 25000 bp) | 918,080,154 |
| Total length (>= 50000 bp) | 916,877,030 |
| # contigs | 149 |
| Largest contig | 44,258,981 |
| Total length | 918,200,210 |
| GC (%) | 52.37 |
| N50 | 15,450,383 |
| N90 | 5,031,844 |
| auN | 16,918,998.2 |
| L50 | 20 |
| L90 | 61 |
| # N's per 100 kbp | 0.00 |

## Methods

### Biomass acquisition

Male gametophytes were collected in Roscoff, France in May 2025 (48°43’47’’N; 4°00’42’’W), under permit No. 98/2025, issued to the Roscoff Biological Station by the Préfet de la Région Bretagne (Direction interrégionale de la mer Nord Atlantique, Manche Ouest, France). Specimens were cultured at the Phycology Research Group at Ghent University. All individuals were properly cleaned by rinsing in several baths of sterile seawater (31 psu) and epiphytes were removed with blotting paper from the thalli. Cultures were grown in filtered seawater (22 µm) at 31 psu, with 10 mL L^-1^ mPES (modified Provasoli Enriched Seawater)^34^ at 12 °C, 12:12 h light:dark photoperiod, and 30 µmol photons m² s^-1^ cool white light.

### DNA extraction, quality control, and library preparation

Genomic DNA was obtained from a single haploid male gametophyte. Before DNA extraction, the biomass was exposed to a series of antibiotics (ABs), as shown in Table 2.

**Table 2.** Scheme of antibiotic treatments before genomic DNA extraction.

| **Day 1** | **Day 2** | **Day 3** | **Day 4** |
|---|---|---|---|
| Streptomycin 0.05 g L<sup>-1</sup> + Penicillin G 0.5 g L<sup>-1</sup> | Ampicillin 0.5 g L<sup>-1</sup> | Cefotaxime 0.5 g L<sup>-1</sup> | Harvest and sonication (5 min at 40 kHz) |

After each AB treatment, the tissue was collected on a 100 µm filter, rinsed 3 times and vortexed with natural seawater (NSW). Finally, tissue was dried using clean blotting paper, and approximately 100 mg of tissue was spread over 10 aliquots in 2 mL Eppendorf tubes and flash frozen using liquid nitrogen. The aliquots were stored at −80°C until DNA extraction

Genomic DNA was extracted using the OmniPrep Genomic DNA Purification Kit (G- Biosciences, USA) according to the manufacturer’s instructions. Extracted DNA was preserved in 100 µl TE buffer and stored at −20 °C until sequencing.

### Nanopore library preparation, quality control, and sequencing

DNA integrity, concentration, and purity were assessed for five aliquots using a Fragment Analyzer (Agilent Technologies, USA), Qubit fluorometer (Thermo Fisher Scientific, USA), and Little Lunatic UV/VIS reader (Unchained Labs, USA). The five independent DNA aliquots yielded between 0.7 and 1.5 µg of DNA with appropriate purity and concentration and an average fragment size of 38-42 kb. Following quality control, all aliquots were pooled irrespective of individual DNA yield to maximize the total amount of input DNA. The pooled DNA was fragmented to an average size of 17–20 kb using the Megaruptor 3 system (Diagenode, Belgium) at speed setting 30. Short DNA fragments were subsequently depleted using the ShortRead Eliminator XS kit (PacBio, USA) according to the manufacturer’s instructions.

Sequencing libraries were prepared using the Oxford Nanopore Technologies SQK-LSK114- XL kit according to the DNA-XL V14 protocol (protocol version GDX_9163_v114_revT_30Jan2025). A total of 35 fmol of the final library was loaded onto a PromethION P24 instrument (Oxford Nanopore Technologies, UK) using a FLO-PRO114M flow cell containing 6,955 active pores at the start of sequencing. Sequencing was performed for 80 h. To maximize sequencing yield, the flow cell was flushed and reloaded every approximate 12 h with an additional 35 fmol of previously prepared library using an in-house optimized reload protocol, resulting in four reloads during the sequencing run. The ONT- sequencing run generated a total 30.34 Gb of data comprising 3.56 million reads, with an N50 read length of 17 kb. Basecalling was performed using the high-accuracy model in MinKNOW v25.03.7 with Dorado v7.8.3 (Oxford Nanopore Technologies, UK), achieving a read pass rate of 98.3%. No quality filters or cut-offs were applied on the base-called data going into subsequent analysis.

### PacBio library preparation, quality control, and sequencing

DNA concentration in the remaining 5 aliquots was determined with Qubit dsDNA High Sensitivity assay (Thermo Fisher Scientific, USA). Integrity was assessed using Fragment Analyzer High Sensitivity Large Fragment kit (Agilent, USA). DNA was size-selected using the PacBio SRE XS kit and converted into SMRTbells following the PacBio “Preparing whole genome and metagenome libraries using SMRTbell^®^ prep kit 3.0” protocol (102-166-600 REV07 JUN2025; PacBio, USA). The resulting SMRTbell was then complexated and sequenced following SMRTlink instructions (SMRT Link: 25.1.0.257715) in one run on the PacBio Revio by the VIB Nucleomics Core (https://nucleomicscore.sites.vib.be/en). Raw sequencing data was demultiplexed and converted to HiFi reads FastQ files using the SMRTLink pipeline and standard PacBio parameters, generating a total Hifi yield of 7.7 Gb raw data.

### RNA-extraction and sequencing

For the integration of RNA-evidence into the BRAKER3 annotation pipeline, mRNA from six individuals that were exposed to different cultivation treatments in the Phycology Research Group Laboratory, was pooled and extracted using the RNeasy^®^ Plant Mini Kit (Qiagen, Hilden, Germany), following the manufacturer’s protocol. These specific individuals, exposed to different environmental parameters (light intensity, salinity, geographic origin), were selected to maximize transcript diversity and support the completeness of the genome annotation. Quality was checked using a NanoDrop™ 2000 spectrophotometer (Thermo Fisher Scientific, USA) and a 5200 Fragment Analyzer using the RNA Kit (15 nt) (Agilent Technologies, USA). Library preparation and high-throughput sequencing were performed by Novogene Co., Ltd. (Cambridge, UK). Quantified libraries were sequenced on the Illumina NovaSeq 6000 platform (PE150), generating an average of 6 Gb of clean reads per sample.

### Flow cytometry

To obtain an estimate of the expected genome size, flow cytometry was performed on both gametophytes and tetrasporophytes from individuals of the same population as the selected reference genome. Prior to nuclei isolation, we subjected the biomass to repeated mechanical cleaning, to remove surface contaminants and associated microbial communities^14^. Young, fresh tips were placed in a 1.5 mL tube containing 1 mL sterile seawater with silica sand ( 50-70 mesh particle size) and were disrupted for 45 s at 25 Hz using a Retsch MM400 mixer mill. This process was repeated two more times, transferring the tissue into a new tube with fresh sterile seawater and silica gel before each round. To finalize the cleaning process, the tissue was rinsed with Milli-Q water and gently blotted dry. For nuclei isolation of *Furcellaria* and our reference control, the clean *Furcellaria* tissue and approximately 20 mg of *Solanum lycopersicum* (cultivar Stupické polní tyčkové rané^35,36^) leaf tissue were co-chopped in ice-cold lysis buffer LB01 using a sterile razorblade. An additional 500 µL of LB01 buffer was added, and the homogenate was stored on ice for several minutes. The solution was then filtered through a 42 µm nylon mesh into a fresh tube and incubated at 4 °C for approximately 1 h during which the sample split into two phases: a lower coloured layer containing remaining cell debris and pigments, and a colourless upper layer enriched in nuclei. 200 μL of this upper layer was isolated and added to a staining solution composed of 550 μL of LB01 lysis buffer, 50 μg mL^−1^ PI, 50 μg mL^−1^ RNase IIA, and 2 μL mL^−1^ β-mercaptoethanol. Following a 10-min dark incubation at 4 °C, the stained samples were analysed with an Attune NxT flow cytometer (Thermo Fisher Scientific, Waltham, MA, USA). Nuclei were identified by using a yellow excitation laser (561 nm, 50 mW, emission filter 620/15, and channel YL2) and subsequently plotting side scatter area against PI intensity. After gating, only the G1-phase nuclei population was retained for analysis, and its median fluorescence intensity was used to estimate genome size^37^.

### *De novo* assembly

We performed a hybrid assembly, with both ONT and PacBio Hifi long reads, using Hifiasm (v0.25.0-r726). To support the assembly of the homozygous gametophytic genome, the *-l0* flag was used to disable the purge duplication step in the Hifiasm pipeline. The resulting *de novo* assembly of *Furcellaria* was 1.00 Gb in length, consisting of 1748 contigs, with a contig N50 of 14.5 Mb and the longest contig consisting of 44.3 Mb, indicating a high level of contiguity and continuity. Initial completeness was evaluated using BUSCO (v6.0.0) against both the eukaryota_odb10 and rhodophyta_odb12 reference databases, revealing 75.6% and 96.5% complete BUSCOs, respectively. Merqury was used to assess the assembly using kmer spectra (k = 21), resulting in a kmer completeness of 99.7% and a consensus quality value (QV) of 30.2. The cumulative contig length plot revealed that most of the assembled genome was represented by a relatively small number of large contigs, whereas the remaining small contigs accounted for only a minor fraction of the total assembly length (Figure 3).

**Figure 3.**
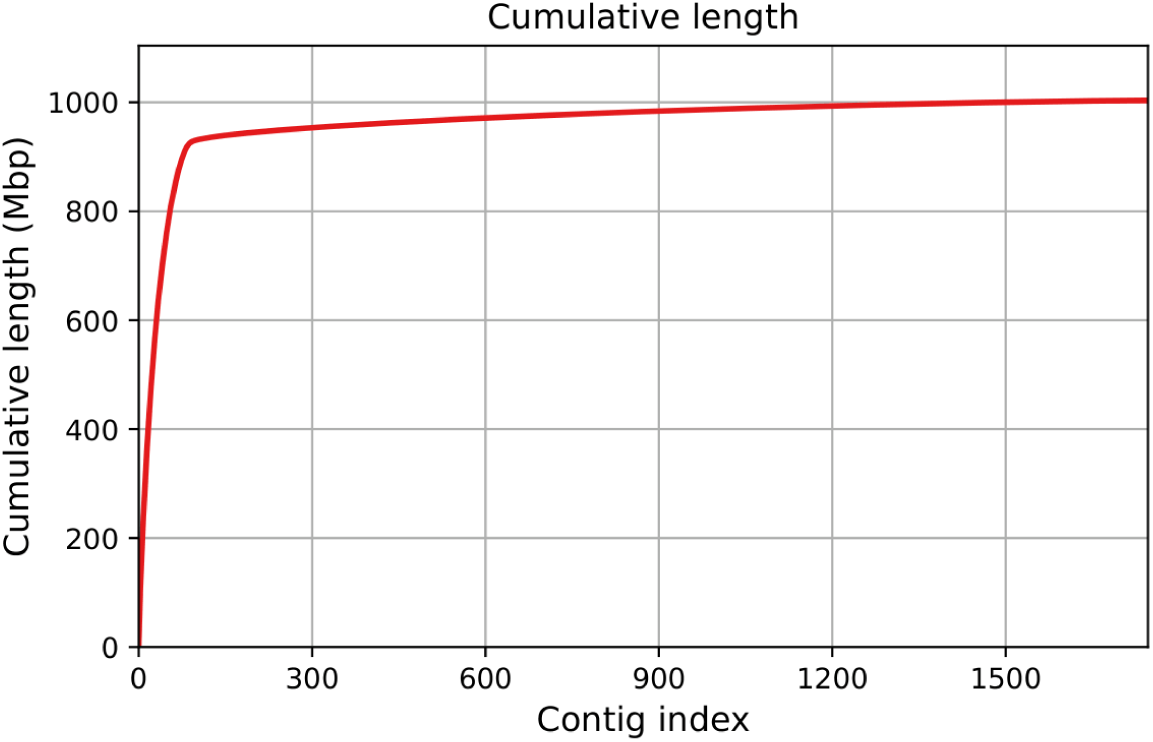
QUAST cumulative contig length plot of the initial *Furcellaria* genome assembly.

### Quality assessment and polishing

To polish the first Hifiasm draft genome, Blobtools (v4.4.5) was used to eliminate all contigs assigned to either *Pseudomonadota* or *Bacterioidea*. This reduced the number of contigs to 949 while maintaining a BUSCO completeness of 76.1 % and a Merqury completeness and QV of 97.6 and 34.0, respectively. The resulting assembly only had a small decrease in size, indicating that mainly numerous small contigs were removed. Additionally, the assembly was further refined by using a hybrid strategy based on both RNA-seq and Illumina DNA read support. RNA-seq data (PE150) from six *Furcellaria* strains representing a European geographic gradient and different cultivation treatments, together with a 40x dataset of PE150 Illumina DNA reads from an individual of the same population, were mapped to the assembly using STAR (v2.7.11b) and BWA-MEM (v0.7.18), respectively. Contigs supported by fewer than five RNA-seq reads and <0.1% of the total Illumina DNA reads (24 million reads) were discarded, yielding a filtered assembly of 451 contigs. BUSCO completeness, Merqury quality values, and genome size remained largely unchanged after this second filtering.

In a final filtering step, TIARA (v1.0.3) was used to identify contigs that were assigned either to mitochondrial or plastid organelles, purging an additional 302 contigs considered non- nuclear. The final, nuclear genome assembly consisted of 149 contigs. BUSCO was performed on the highest related taxonomic level (eukaryota_odb10), resulting in a score of 74.9% and a lower taxonomic level (rhodophyta_odb12) resulting in a significantly higher BUSCO of 96.7%, indicating the completeness within the Rhodophyta genomes assembled to date. Merqury kmer (k=21) completeness and QV-scores were 93.17 and 40.7, respectively.

### Repeat annotation and soft-masking of genome

Before predicting protein-coding genes in the genome, repeat annotation was conducted using a combination of *de novo* and homology-based approaches to identify and mask repetitive elements in the genome. We first used RepeatModeler (v.2.0.5) to build a *de novo* custom-built and species-specific library generated for *Furcellaria* to increase the accuracy of detection and annotation of transposable elements (TEs). We used the NCBI BLAST engine for sequence comparisons during repeat classification. The resulting custom repeat library was then given as input for RepeatMasker (v4.1.5) (RM) using the Dfam database for identification and soft masking (*cutoff score = 225, -xmall, -gff, -excln*) of TEs in the *Furcellaria* genome. To resolve tandem repeats, TandemRepeatFinder (v4.09.1) (TRF) was used with the following scoring parameters; *match = 2, mismatch = 7, delta =7, PM = 80, PI = 10, minscore = 50, maxperiod = 2000.* The resulting coordinates of the regions identified by RepeatMasker and TRF were combined using BEDtools (v2.31.1) with the *-maskfasta* option to soft-mask all identified repeats in the final genome assembly. This two-fold strategy identified a substantial number of repetitive elements, dominated by DNA transposons and retroelements. The total masked fraction comprises 90.5% of the genome (Table 3; Supplementary Information Figure S1).

**Table 3.** Overview of repeat content and annotation in the *Furcellaria* genome assembly.

| Repeatclass/family | Occupied length (bp) | % of genome |
|---|---|---|
| **Interspersed repeats (RM)** |  |  |
| **Retroelements (Class I)** | 310,099,568 | 33.77 |
| --LTR elements | 288,126,184 | 31.38 |
| ------Gypsy/DIRS1 | 170,179,316 | 18.53 |
| ------BEL/Pao | 73,400,219 | 7.99 |
| ------Ty1/Copia | 43,015,266 | 4.68 |
| --Penelope | 174,337 | 0.02 |
| --LINEs | 21,715,122 | 2.36 |
| --SINEs | 258,262 | 0.03 |
| **DNA transposons (Class II)** | 355,668,228 | 38.74 |
| **Rolling-circle transposons** | 3,626,620 | 0.39 |
| **Unclassified repeats** | 151,706,053 | 16.52 |
| **Other repeats (simple repeats, small RNA, low complexity)** | 5,503,768 | 0.60 |
| **Tandem repeats (TRF)** |  |  |
| Additional tandem repeats | 4,596,034 | 0.5 |
| **TOTAL MASKED FRACTION (RM + TRF)** | 831,116,346 | **90.5** |

### Structural genome annotation

Adaptor-trimmed RNA-seq reads were aligned to the soft-masked *Furcellaria* genome using STAR (v2.7.11b) for optimal discovery of novel splice junctions and intron evidence, with the *--outSAMstrandField intronMotif* option to retain strand information for spliced alignments. Given the repetitive nature of the genome, secondary alignments were removed prior to BRAKER3 gene prediction, resulting in 353 million primary alignments that were used as transcript evidence^38^. Additionally, a curated set of protein orthologues was derived from ten published Rhodophyta proteomes (Table 4). Orthologues were first identified using OrthoFinder (v2.5.5), yielding an initial 9187 orthogroups. This dataset was subsequently filtered to obtain only those orthogroups present in at least 80% (8) of the reference species, resulting in a final dataset of 3962 conserved orthogroups, merged into a single database to serve as protein homology evidence for subsequent BRAKER3 gene prediction.

**Table 4.**
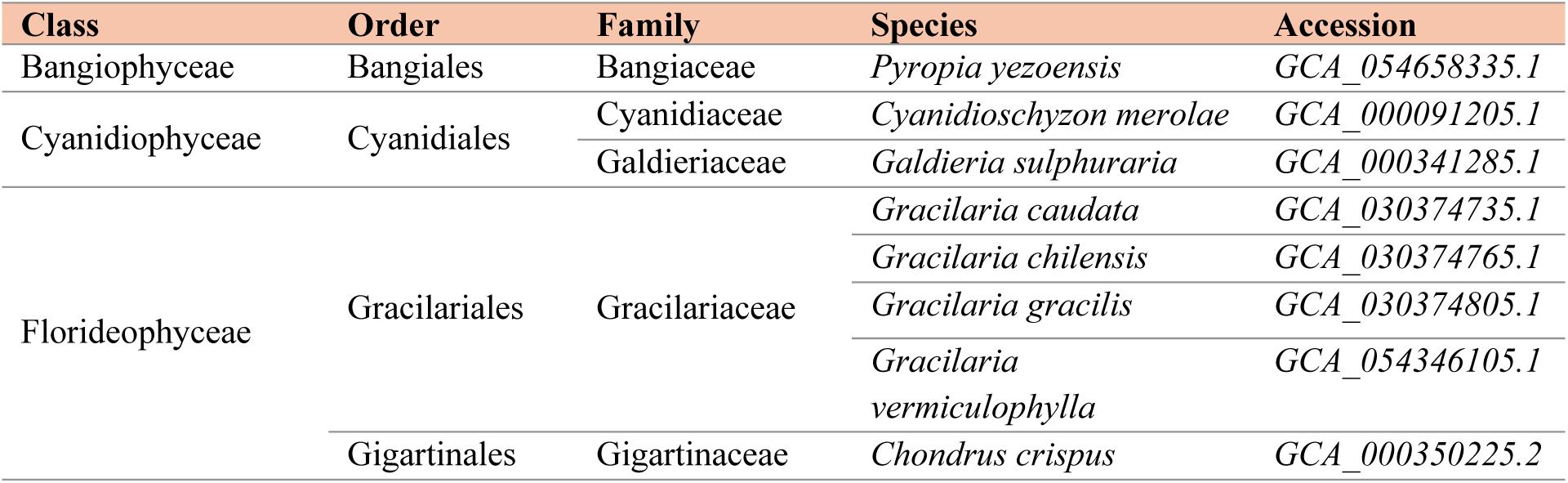
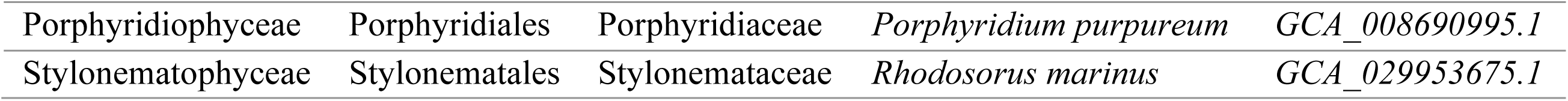
Overview of Rhodophyta used for protein homology evidence.

The soft-masked genome was used as input for BRAKER3 (v3.0.8), together with the generated RNA-seq evidence and homologous protein info, resulting in a single GFF3 annotation file. Providing both RNA and protein evidence, BRAKER3 was executed with GeneMark-ETP. All other parameters were kept default. To refine the initial output, AGAT (v1.4.0) was used for the purging of redundant isoforms. Structural annotation statistics are summarized in Table 5.

**Table 5.** Gene and annotation features after isoform-purging of the newly assembled *Furcellaria* genome.

| **Feature** | **Value** |
|---|---|
| Protein-coding genes | 6100 |
| mRNAs | 6100 |
| CDs | 6100 |
| Exons | 8124 |
| Introns | 2024 |
| Mean gene length | 1684 |
| Mean CDS length | 1301 |
| Mean exons per gene | 1.3 |
| Mean intron length | 1156 |
| % genes with functional annotation | 82.6% (5037) |
| % genes with GO terms | 73.9% (4509) |
| % genes with InterPro domains | 80.0% (4876) |

### Functional genome annotation

Functional annotation was performed using both eggNOG-mapper (v2.1.13) and InterProScan (v5.59-91.0). EggNOG-mapper provided orthology-based functional assignments, including Gene Ontology (GO) terms, KEGG pathways, and COG categories, while InterProScan identified conserved protein domains and domain-derived GO terms (Figure 4, Figure 5). GO terms were combined to maximize functional coverage. Functional annotation statistics are shown in Table 5.

**Figure 4.**
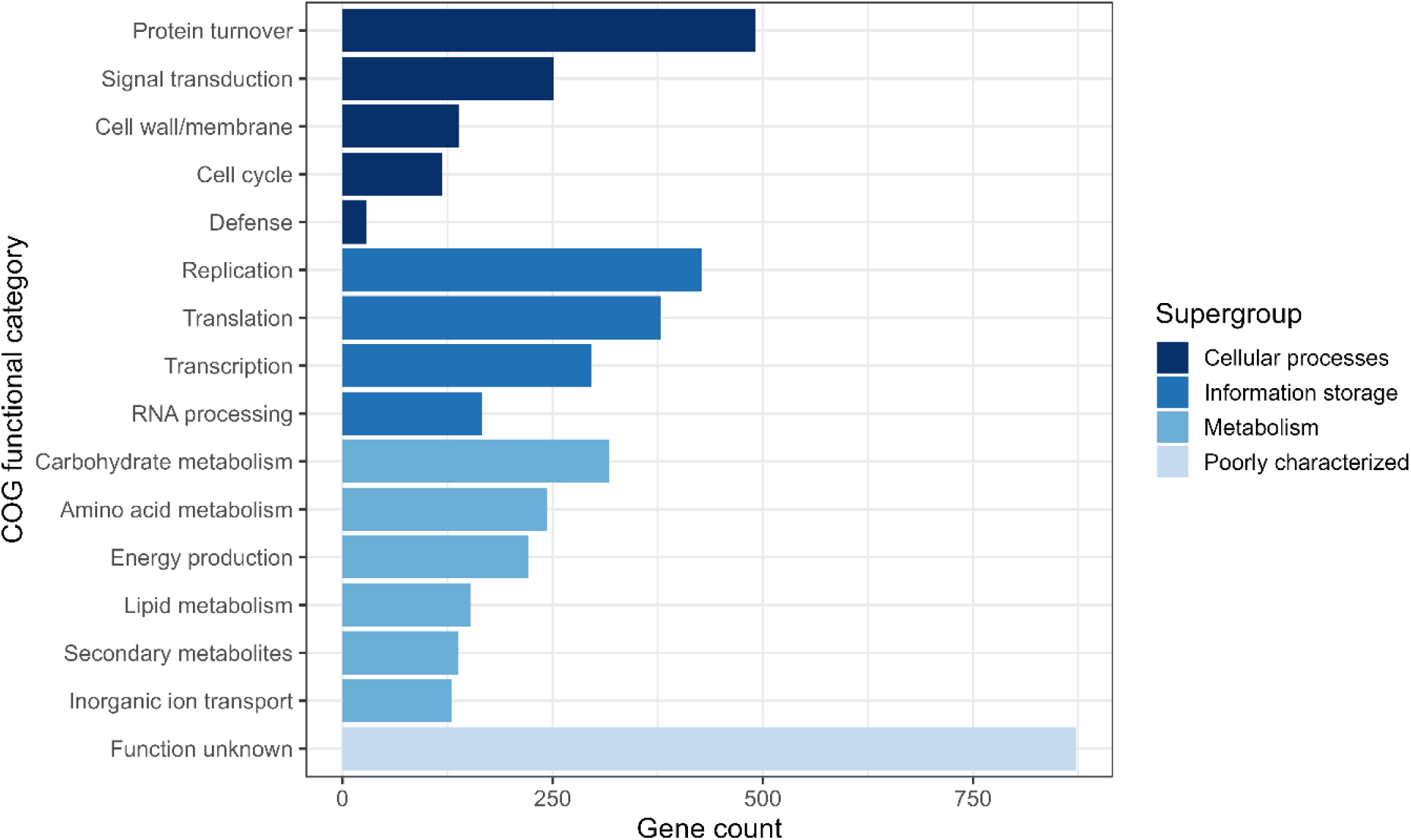
Functional classification of predicted protein-coding genes in *Furcellaria* based on COG-annotations assigned by eggNOG-mapper. Bars indicate the number of genes assigned to each major functional category.

**Figure 5.**
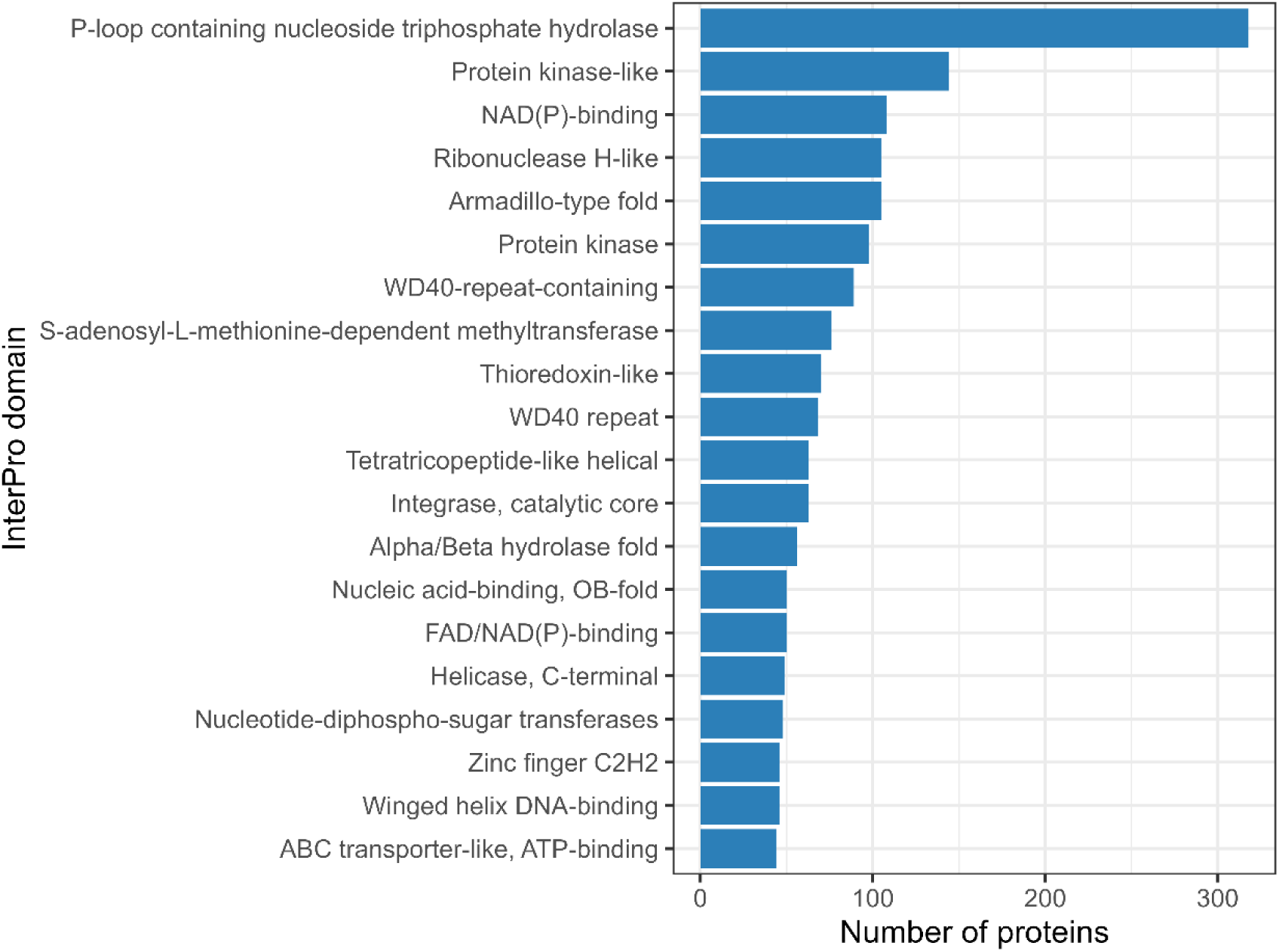
Top 20 most abundant InterPro protein domains identified in the predicted proteome of *Furcellaria*. Domain assignments were obtained using InterProScan and bars indicate the number of proteins containing each domain.

### Mitochondrial and chloroplast genome assemblies

Organellar scaffolds, as identified by TIARA (v1.0.3) were extracted and used for subsequent assembly of the mitochondrial and chloroplast genomes. For assembling the mitochondrial genome, MitoHifi (v3.2.3) was used with the reference mitogenome of *Gibsmithia punonomaewa* (PV035080) as the closest relative. This resulted in the assembly of a 31.5 Kb mitochondrial genome. The chloroplast genome was assembled and using NOVOplasty (v4.3.5), resulting in a 187,5 Kb long genome.

Genes were initially identified by mapping the genes from the *Chondrus crispus* mitochondrial (NC_001677.2) and chloroplast (HF562234) genomes against the *Furcellaria* organellar contigs using the ‘Map to Reference’ tool with high-sensitivity parameters in Geneious Prime 2026.0.2. Annotation completeness was assessed using the ‘Find ORFs’ tool in Geneious, followed by BLASTp searches against the NCBI nonredundant database. tRNA genes were verified using tRNAscan-SE 2.0. A graphic overview of the organellar assemblies is provided in the Supplementary Information Figure S2.

## Data records

The PacBio HiFi and ONT long read data have been deposited in the Sequence Read Archive (SRA) with accession numbers SRR37327885 and SRR37327886, respectively, under BioProject PRJNA1424984. The genome assembly has been submitted to NCBI GenBank under the accession JBVBPQ000000000. Genome annotation files are available under https://doi.org/10.6084/m9.figshare.33093305. mtDNA and cpDNA and their annotation features have been submitted to NCBI GenBank under the accessions PZ734586 and PZ720091, respectively. All data corresponding to the *Furcellaria* reference genome assembly is linked to BioSample SAMN55401920.

## Technical validation

Flow cytometry was performed to estimate the genome size of *Furcellaria* and to assess whether the flow cytometry derived estimates coincided with the genome size obtained in the final genome assembly. *Solanum lycopersicum* (cultivar Stupické polní tyčkové rané), with a 2C DNA content of 1.96 pg, was used as an internal reference^39,40^. For completeness, both a male gametophyte (1C) and a tetrasporophyte (2C) were included, yielding estimated genome size intervals of 920–980 Mb and 1900–1950 Mb, respectively (Figure 6). These estimates approximate the haploid assembly span, further supporting the completeness of our assembled reference.

**Figure 6.**
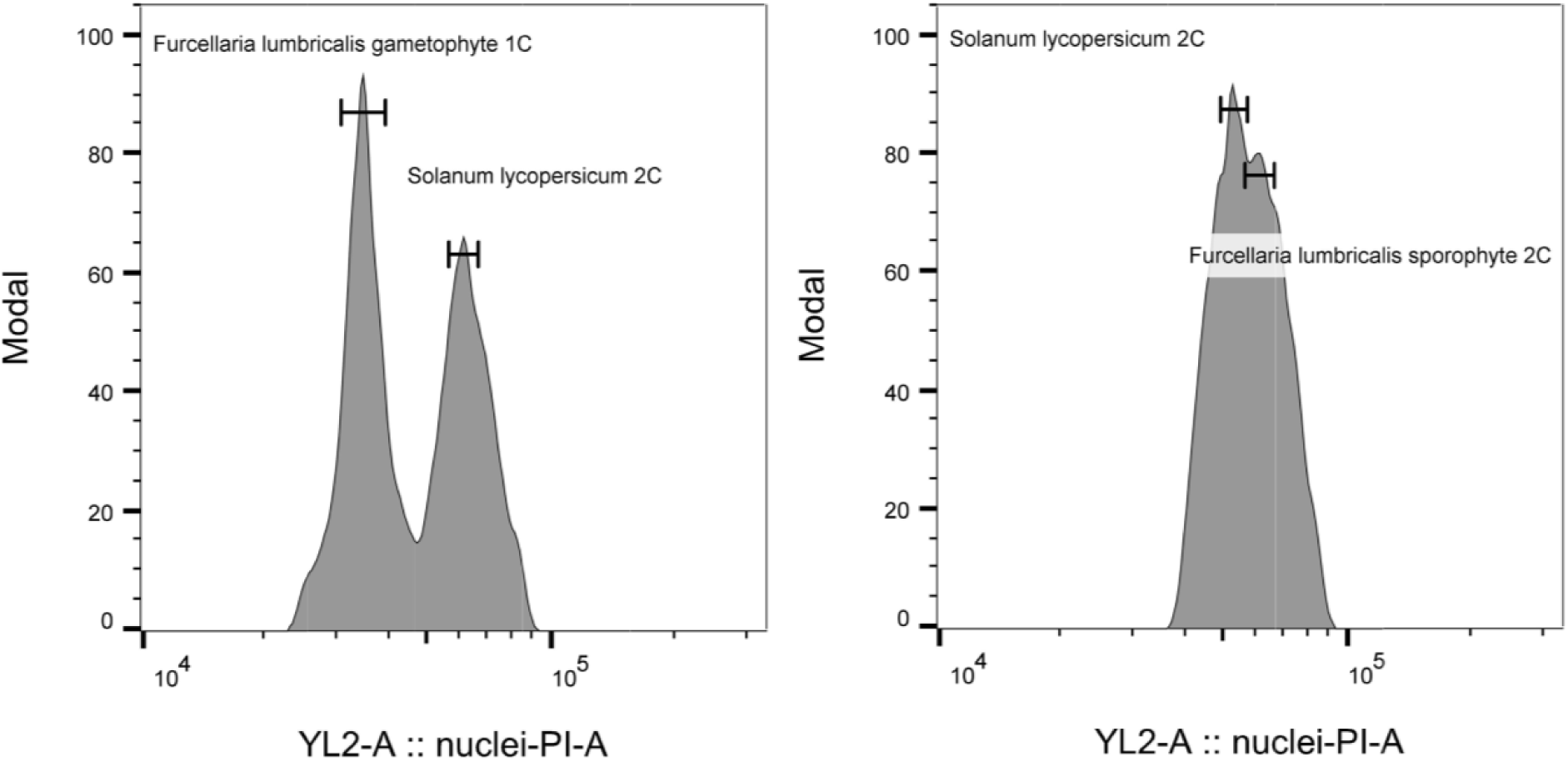
Flow cytometry profiles of propidium iodide (PI)-stained nuclei, gated for the G1-phase of the cell cycle in each species. Left: mixed analysis of gametophytic biomass (n) of *Furcellaria* with leaf tissue of *Solanum lycopersicum*, right: mixed analysis of tetrasporophyte biomass (2n) of *Furcellaria* with leaf tissue of *Solanum lycopersicum*.

Completeness and quality of the *Furcellaria* genome were assessed throughout the assembly and polishing process, using QUAST, BUSCO, Merqury, BlobTools, and TIARA (sections: *de novo* assembly and polishing). To further evaluate the assembly and the representation of the gene space, Illumina genomic DNA and RNA-seq reads were re-mapped to the final polished assembly using BWA-MEM and STAR, respectively. Overall alignment rates exceeded 90%, with average mapping quality scores exceeding 30.

## Code availability

All software and pipelines were executed according to the manual, GitHub repositories, and protocols of the cited software packages and tools (Table 6). Unless otherwise specified, all tools were executed using default parameter settings as described in the official documentation. Any additional information or specific command-line implementations required to reanalyse the data reported in this study are available from the corresponding author upon request.

**Tabel 6:**
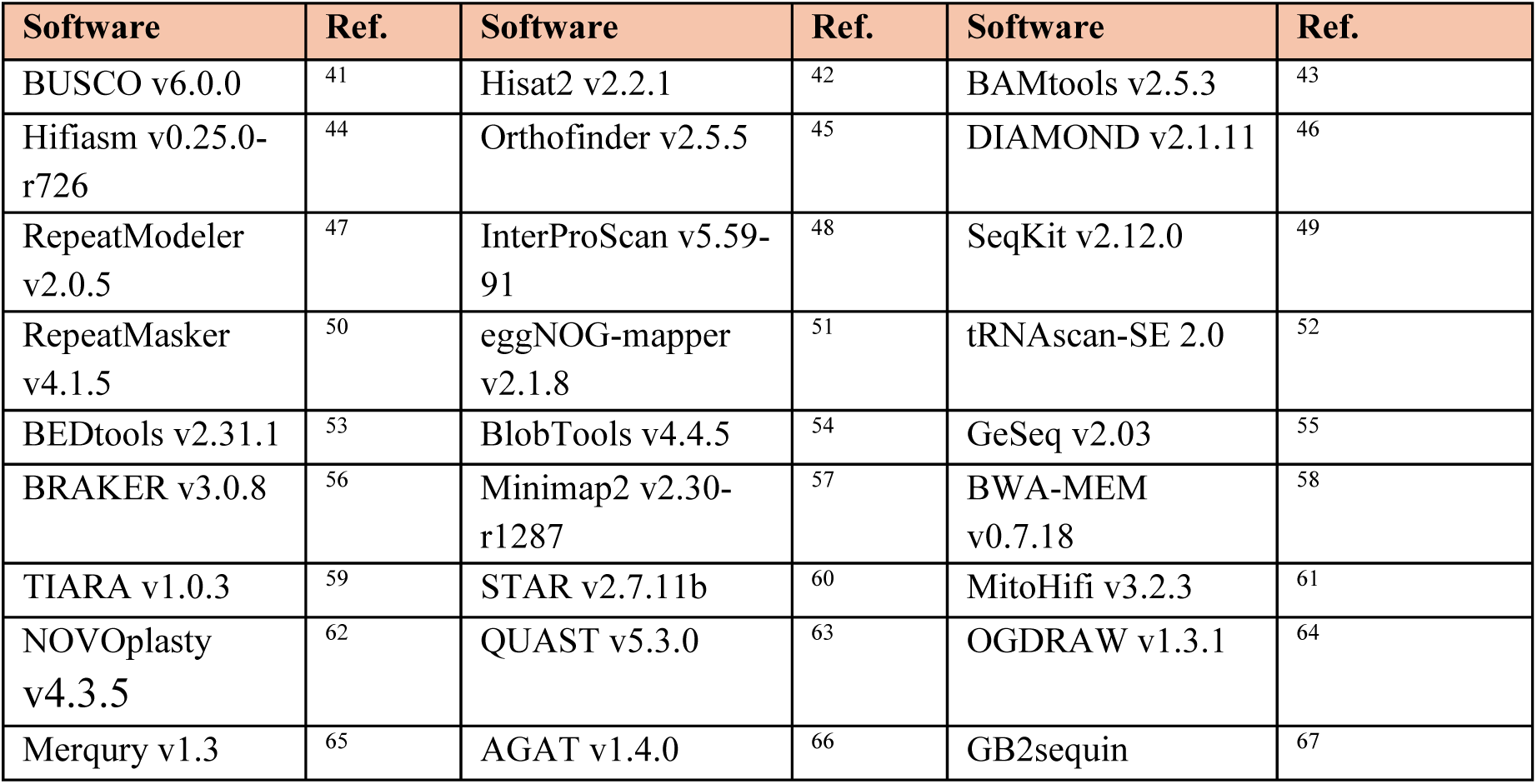
Overview of used software in the genome assembly pipeline.

## Author Contributions

**O.D.C.**, **J.K.**, and **K.A.** secured project funding and provided overall project supervision and facilitation. **L.N.** collected all biological samples and was responsible for the algal cultures. **S.D.** and **L.N.** were involved in optimization of DNA extraction, and QC. **M.S.** and **T.D.P.** conceptualised ONT sequencing experiment, performed quality control, ONT sample and library preparation, sequencing, reloading and helped with writing of the ONT-sections in the manuscript. **L.N.** performed all genome assemblies, polishing, and annotations, with additional bioinformatic support from **J.M.** and **A.V.d.V. L.N.** performed technical validation, with help from **A.V.d.V.** for generating flow cytometry genome size estimates. **F.L.** provided support in the annotation of the organellar genomes, writing, as well as the NCBI submissions. **L.N.** managed data curation and submission onto public repositories. **L.N.** drafted the first version of the manuscript. All authors reviewed, edited, and approved the final version of the manuscript.

## Data Availability Statement

All data have been submitted to the corresponding repositories, as described in the ‘Data Records’ section, and will become publicly available upon publication of the manuscript under their respective accession numbers. Raw sequence data (Hifi + ONT) has been deposited onto SRA and will become available upon publication via the accession numbers as described in ‘Data Records’.

## Acknowledgements

We thank Station Biologique de Roscoff, especially Ronan Garnier, for the support in collecting the algae and the shipment to Belgium. We further thank the Neuromics Support Facility (VIB-UAntwerp Center for Molecular Neurology) and the Nucleomics Core Facility, both part of the Vlaams Instituut voor Biotechnologie (VIB), for long-read sequencing and data provision. Finally, we would like to thank Quinten Bafort (Ghent University, Phycology research Group and Department of Plant Biotechnology and Bioinformatics), Romy Petroll and Michael Borg (both from the Max Planck Institute for Biology, Tübingen, Germany), for all their support and guidance during the assembly stages of the nuclear genome.

---

**Figure S1.**
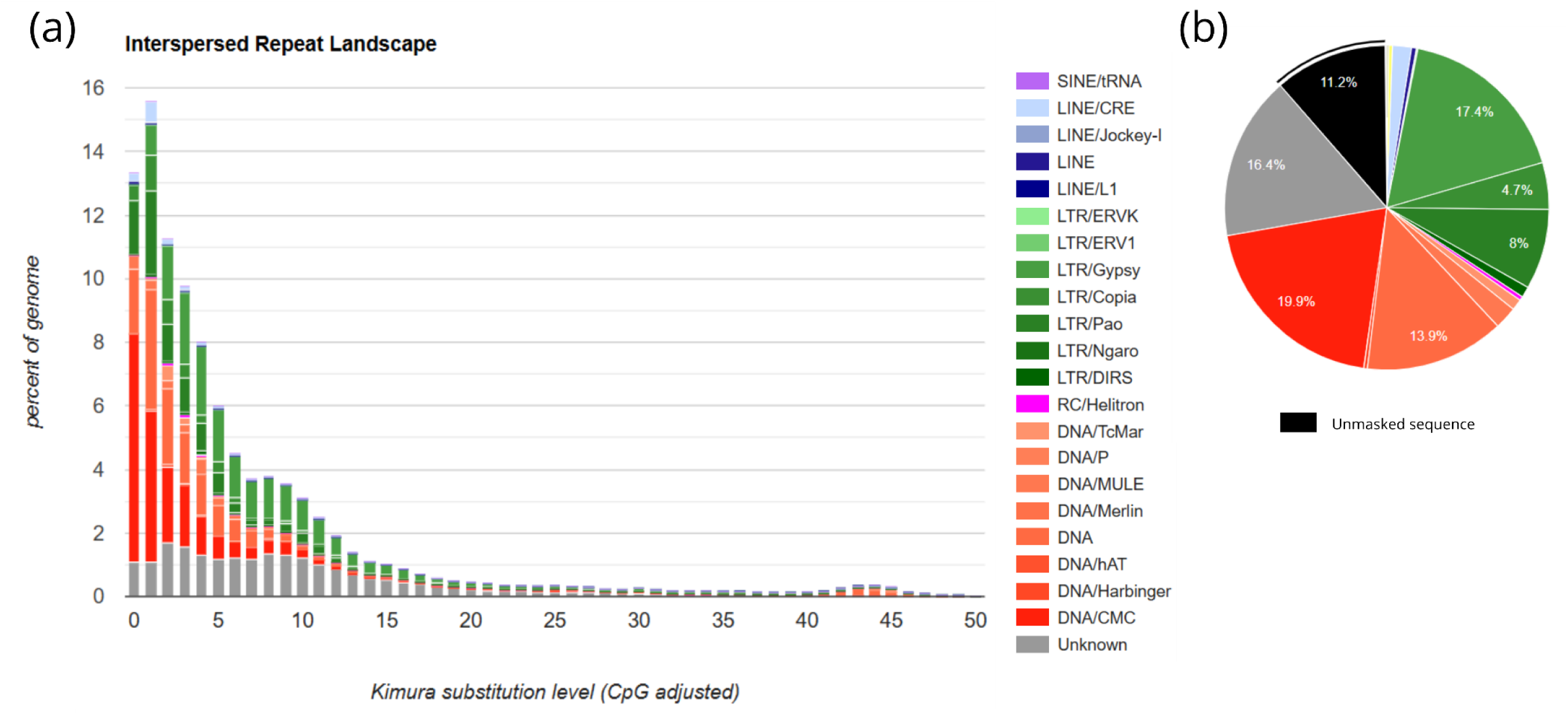
Repeat landscape and composition of repetitive elements in the *Furcellaria* genome. **(a)** Repeat landscape showing the genomic proportion occupied by major repeat classes across Kimura substitution levels (CpG-adjusted), where lower divergence values indicate more recent repeat insertions. The landscape is dominated by DNA transposons and LTR retrotransposons, with a pronounced peak at low divergence values, indicating recent repeat emergence. **(b)** Genomic fraction occupied by the major repeat classes identified by RepeatMasker.

**Figure S2.**
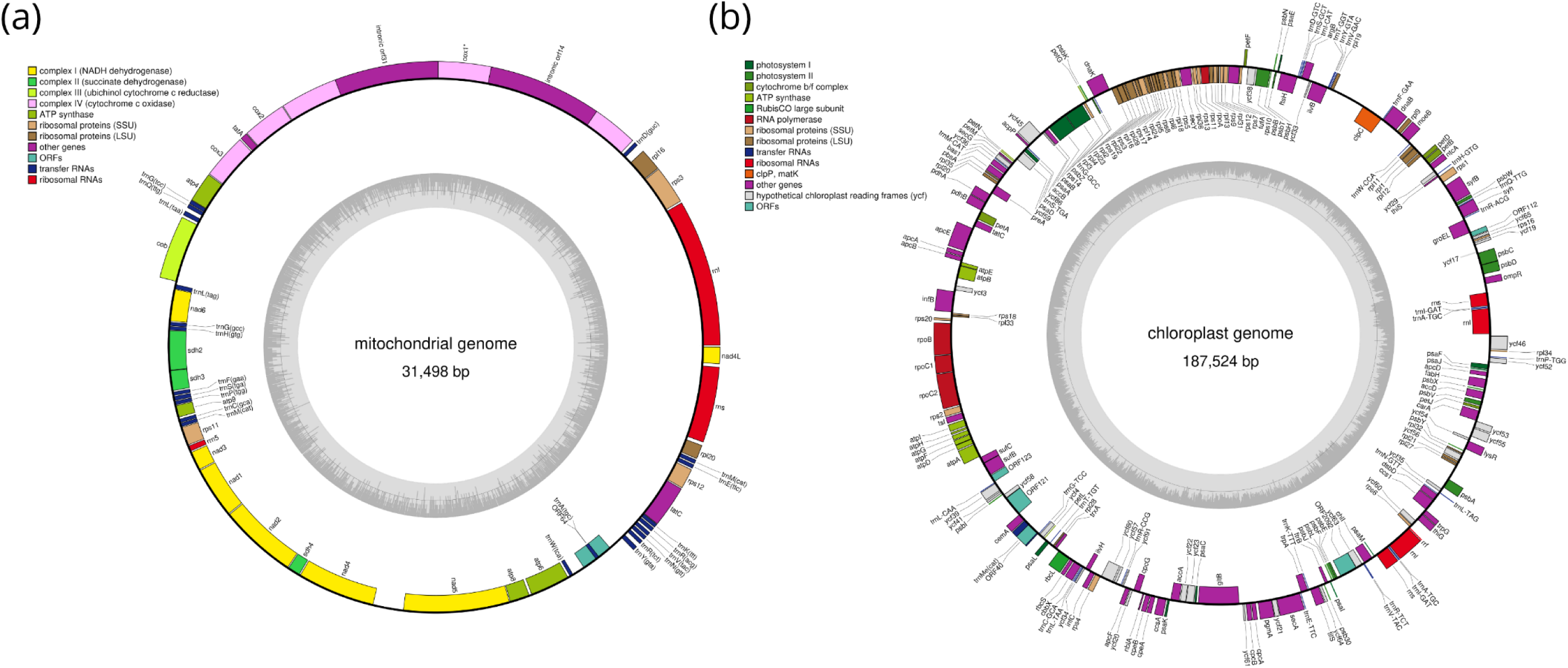
Visual representation of the annotation of the newly assembled mitochondrial (a) and chloroplast (b) genome of *Furcellaria*. Genome maps drawn with OGDRAW^64^.

## References

1. Ruiz-Trillo, I. & López-Escardó, D. Multicellularity: Red algal genomic expansion fine-tuned with transposable elements. Current Biology 35, R606– R608 (2025).

2. Borg, M. et al. Red macroalgae in the genomic era. New Phytologist 240, 471– 488 (2023).

3. Cock, J. M. & Collén, J. Independent Emergence of Complex Multicellularity in the Brown and Red Algae. in Evolutionary Transitions to Multicellular Life: Principles and mechanisms (eds. Ruiz-Trillo, I. & Nedelcu, A. M.) 335–361 (Springer Netherlands, Dordrecht, 2015). doi:10.1007/978-94-017-9642-2_16.

4. Pearman, W. S. et al. A cry for kelp: Evidence for polyphenolic inhibition of Oxford Nanopore sequencing of brown algae. J. Phycol. 60, 1601–1610 (2024).

5. Vidal, R., Meneses, I. & Smith, M. Enhanced DNA extraction and PCR amplification of SSU ribosomal genes from crustose coralline algae. J. Appl. Phycol. 14, 223–227 (2002).

6. Lipinska, A. P. et al. The Rhodoexplorer Platform for Red Algal Genomics and Whole-Genome Assemblies for Several Gracilaria Species. Genome Biol. Evol. 15, evad124 (2023).

7. Jian, J. et al. Chromosome-level genome assembly of agar-producing red seaweed Gracilaria vermiculophylla. Sci. Data 13, 334 (2026).

8. Brawley, S. H. et al. Insights into the red algae and eukaryotic evolution from the genome of Porphyra umbilicalis (Bangiophyceae, Rhodophyta). Proceedings of the National Academy of Sciences 114, E6361–E6370 (2017).

9. Cao, M. et al. A chromosome-level genome assembly of Pyropia haitanensis (Bangiales, Rhodophyta). Mol. Ecol. Resour. 20, 216–227 (2020).

10. Wang, D. et al. Pyropia yezoensis genome reveals diverse mechanisms of carbon acquisition in the intertidal environment. Nat. Commun. 11, 4028 (2020).

11. Nakamura, Y. et al. The First Symbiont-Free Genome Sequence of Marine Red Alga, Susabi-nori (Pyropia yezoensis). PLoS One 8, e57122- (2013).

12. Appeltans, W. et al. The Magnitude of Global Marine Species Diversity. Current Biology 22, 2189–2202 (2012).

13. Kapraun, D. F. & Freshwater, D. W. Estimates of nuclear DNA content in red algal lineages. AoB Plants 2012, pls005 (2012).

14. Petroll, R. et al. The expanded Bostrychia moritziana genome unveils evolution in the most diverse and complex order of red algae. Current Biology 35, 2771–2788.e8 (2025).

15. Qiu, H., Price, D. C., Yang, E. C., Yoon, H. S. & Bhattacharya, D. Evidence of ancient genome reduction in red algae (Rhodophyta). J. Phycol. 51, 624–636 (2015).

16. El Gamal, A. A. Biological importance of marine algae. Saudi Pharmaceutical Journal 18, 1–25 (2010).

17. Kersen, P., Paalme, T., Pajusalu, L. & Martin, G. Biotechnological applications of the red alga Furcellaria lumbricalis and its cultivation potential in the Baltic Sea. 60, 207–218 (2017).

18. Leal, M. C. et al. Biogeography and biodiscovery hotspots of macroalgal marine natural products. Nat. Prod. Rep. 30, 1380–1390 (2013).

19. Flores-Molina, M. R. et al. Desiccation stress in intertidal seaweeds: Effects on morphology, antioxidant responses and photosynthetic performance. Aquat. Bot. 113, 90–99 (2014).

20. Davison, I. R. & Pearson, G. A. STRESS TOLERANCE IN INTERTIDAL SEAWEEDS. J. Phycol. 32, 197–211 (1996).

21. Dagnino-Leone, J. et al. Phycobiliproteins: Structural aspects, functional characteristics, and biotechnological perspectives. Comput. Struct. Biotechnol. J. 20, 1506–1527 (2022).

22. Gaspar, R., Pereira, L. & Neto, J. M. Intertidal zonation and latitudinal gradients on macroalgal assemblages: Species, functional groups and thallus morphology approaches. Ecol. Indic. 81, 90–103 (2017).

23. Bird, C. J., Saunders, G. W. & McLachlan, J. Biology of Furcellaria lumbricalis (Hudson) Lamouroux (Rhodophyta: Gigartinales), a commercial carrageenophyte. J. Appl. Phycol. 3, 61–82 (1991).

24. Bird, C. J., Saunders, G. W. & McLachlan, J. Biology of Furcellaria lumbricalis (Hudson) Lamouroux (Rhodophyta: Gigartinales), a commercial carrageenophyte. J. Appl. Phycol. 3, 61–82 (1991).

25. Weinberger, F., Paalme, T. & Wikström, S. A. Seaweed resources of the Baltic Sea, Kattegat and German and Danish North Sea coasts. 63, 61–72 (2020).

26. Kotta, J. et al. Density dependent growth of the red algae Furcellaria lumbricalis and Coccotylus truncatus in the West Estonian Archipelago Sea, northern Baltic Sea. Oceanologia 50, (2008).

27. Rinne, H. & Kostamo, K. Distribution and species composition of red algal communities in the northern Baltic Sea. Estuar. Coast. Shelf Sci. 269, 107806 (2022).

28. Skapste, I., Grinberga-Zalite, G. & Žaimis, U. Economic Potential of Algae Biostimulant for Sustainable Agriculture in the Baltic Sea Region: Impact of Furcellaria lumbricalis Digestate Extract on Basil Growth Promotion. Sustainability 17, (2025).

29. Romagnoli, F. et al. Furcellaria lumbricalis macroalgae cascade biorefinery: a Life Cycle Assessment study in the Baltic Sea Region. J. Clean. Prod. 478, 143861 (2024).

30. Saluri, M., Kaldmäe, M. & Tuvikene, R. Extraction and quantification of phycobiliproteins from the red alga Furcellaria lumbricalis. Algal Res. 37, 115– 123 (2019).

31. Romagnoli, F. et al. Furcellaria lumbricalis macroalgae cascade biorefinery: a Life Cycle Assessment study in the Baltic Sea Region. J. Clean. Prod. 478, 143861 (2024).

32. Wennerström, L., Jansson, E. & Laikre, L. Baltic Sea genetic biodiversity: Current knowledge relating to conservation management. Aquat. Conserv. 27, 1069–1090 (2017).

33. OBIS (2026). Ocean Biodiversity Information System. Intergovernmental Oceanographic Commission of UNESCO. https://obis.org.

34. West, J. A. & McBride, D. L. Long-term and diurnal carpospore discharge patterns in the Ceramiaceae, Rhodomelaceae and Delesseriaceae (Rhodophyta). Hydrobiologia 398, 101–114 (1999).

35. Dpooležel, J., Sgorbati, S. & Lcretti, S. Comparison of three DNA fluorochromes for flow cytometric estimation of nuclear DNA content in plants. Physiol. Plant. 85, 625–631 (1992).

36. Dpooležel, J., Binarová, P. & Lcretti, S. Analysis of Nuclear DNA content in plant cells by Flow cytometry. Biol. Plant. 31, 113–120 (1989).

37. Sliwinska, E. et al. Application-based guidelines for best practices in plant flow cytometry. Cytometry Part A 101, 749–781 (2022).

38. Raplee, I. D., Evsikov, A. V & de Evsikova, C. Aligning the Aligners: Comparison of RNA Sequencing Data Alignment and Gene Expression Quantification Tools for Clinical Breast Cancer Research. J. Pers. Med. 9, (2019).

39. Soni, A. & Henry, R. J. Re-calibration of flow cytometry standards for plant genome size estimation. Front. Plant Sci. Volume 16–2025, (2025).

40. Doležel, J., Čížková, J., Šimková, H. & Bartoš, J. One Major Challenge of Sequencing Large Plant Genomes Is to Know How Big They Really Are. Int. J. Mol. Sci. 19, (2018).

41. Tegenfeldt, F. et al. OrthoDB and BUSCO update: annotation of orthologs with wider sampling of genomes. Nucleic Acids Res. 53, D516–D522 (2025).

42. Kim, D., Paggi, J. M., Park, C., Bennett, C. & Salzberg, S. L. Graph-based genome alignment and genotyping with HISAT2 and HISAT-genotype. Nat. Biotechnol. 37, 907–915 (2019).

43. Barnett, D. W., Garrison, E. K., Quinlan, A. R., Strömberg, M. P. & Marth, G. T. BamTools: a C++ API and toolkit for analyzing and managing BAM files. Bioinformatics 27, 1691–1692 (2011).

44. Cheng, H., Asri, M., Lucas, J., Koren, S. & Li, H. Scalable telomere-to-telomere assembly for diploid and polyploid genomes with double graph. Nat. Methods 21, 967–970 (2024).

45. Emms, D. & Kelly, S. OrthoFinder: Phylogenetic orthology inference for comparative genomics. Genome Biol. 20, (2019).

46. Buchfink, B., Reuter, K. & Drost, H.-G. Sensitive protein alignments at tree-of- life scale using DIAMOND. Nat. Methods 18, 366–368 (2021).

47. Flynn, J. et al. RepeatModeler2 for automated genomic discovery of transposable element families. Proceedings of the National Academy of Sciences 117, 201921046 (2020).

48. Jones, P. et al. InterProScan 5: genome-scale protein function classification. Bioinformatics 30, 1236–1240 (2014).

49. Shen, W., Sipos, B. & Zhao, L. SeqKit2: A Swiss army knife for sequence and alignment processing. iMeta 3, e191 (2024).

50. Chen, N. Using RepeatMasker to Identify Repetitive Elements in Genomic Sequences. *Current protocols in bioinformatics / editoral board*, Andreas D. Baxevanis … [et al.] **Chapter** 4, Unit 4.10 (2004).

51. Cantalapiedra, C. P., Hernández-Plaza, A., Letunic, I., Bork, P. & Huerta-Cepas, J. eggNOG-mapper v2: Functional Annotation, Orthology Assignments, and Domain Prediction at the Metagenomic Scale. Mol. Biol. Evol. 38, 5825–5829 (2021).

52. Lowe, T. M. & Chan, P. P. tRNAscan-SE On-line: integrating search and context for analysis of transfer RNA genes. Nucleic Acids Res. 44, W54–W57 (2016).

53. Quinlan, A. R. & Hall, I. M. BEDTools: a flexible suite of utilities for comparing genomic features. Bioinformatics 26, 841–842 (2010).

54. Laetsch, D. R. & Blaxter, M. L. BlobTools: Interrogation of genome assemblies [version 1; peer review: 2 approved with reservations] . F1000Res. 6, (2017).

55. Tillich, M. et al. GeSeq – versatile and accurate annotation of organelle genomes. Nucleic Acids Res. 45, W6–W11 (2017).

56. Gabriel, L., et al. BRAKER3: Fully automated genome annotation using RNA- seq and protein evidence with GeneMark-ETP, AUGUSTUS and TSEBRA. bioRxiv 2023.06.10.544449 (2024) doi:10.1101/2023.06.10.544449.

57. Li, H. Minimap2: pairwise alignment for nucleotide sequences. Bioinformatics 34, 3094–3100 (2018).

58. Li, H. & Durbin, R. Fast and accurate short read alignment with Burrows– Wheeler transform. Bioinformatics 25, 1754–1760 (2009).

59. Karlicki, M., Antonowicz, S. & Karnkowska, A. Tiara: deep learning-based classification system for eukaryotic sequences. Bioinformatics 38, 344–350 (2022).

60. Dobin, A. & Gingeras, T. R. Mapping RNA-seq Reads with STAR. Curr. Protoc. Bioinformatics 51, 11.14.1–11.14.19 (2015).

61. Uliano-Silva, M. et al. MitoHiFi: a python pipeline for mitochondrial genome assembly from PacBio high fidelity reads. BMC Bioinformatics 24, 288 (2023).

62. Dierckxsens, N., Mardulyn, P. & Smits, G. NOVOPlasty: de novo assembly of organelle genomes from whole genome data. Nucleic Acids Res. 45, e18–e18 (2017).

63. Gurevich, A., Saveliev, V., Vyahhi, N. & Tesler, G. QUAST: quality assessment tool for genome assemblies. Bioinformatics 29, 1072–1075 (2013).

64. Greiner, S., Lehwark, P. & Bock, R. OrganellarGenomeDRAW (OGDRAW) version 1.3.1: expanded toolkit for the graphical visualization of organellar genomes. Nucleic Acids Res. 47, W59–W64 (2019).

65. Rhie, A., Walenz, B. P., Koren, S. & Phillippy, A. M. Merqury: reference-free quality, completeness, and phasing assessment for genome assemblies. Genome Biol. 21, 245 (2020).

66. 66. Jacques, D. NBISweden/AGAT: AGAT-v0.0.1. Preprint at 10.5281/zenodo.3549547 (2019).

67. Lehwark, P. & Greiner, S. GB2sequin - A file converter preparing custom GenBank files for database submission. Genomics 111, 759–761 (2019).

